# Quantitative Comparison of Monomeric StayGold Variants Using Protein Nanocages in Living Cells

**DOI:** 10.1101/2024.09.16.613379

**Authors:** Giulia Viola, Kyle A. Jacobs, Joël Lemière, Matthew L. Kutys, Torsten Wittmann

**Affiliations:** Department of Cell and Tissue Biology, University of California, San Francisco, San Francisco, CA 94941

## Abstract

To standardize comparison of fluorescent proteins and independently determine which monomeric StayGold variant is best for live microscopy, we analyzed fluorescent protein tagged I3-01 peptides that self-assemble into stable sixty subunit dodecahedrons inside live cells. We find mStayGold is 3-fold brighter and 3-fold more photostable compared with EGFP and superior to other monomeric variants in mammalian cytoplasm. In addition, analysis of intracellular nanocage diffusion confirms the monomeric nature of mStayGold.

---

Fluorescent proteins (FPs) have transformed cell biology by allowing for real-time observation of dynamic biological processes, from the level of entire organisms down to individual proteins. However, photobleaching and phototoxicity pose critical limitations, particularly in high-resolution live microscopy experiments to study intracellular protein dynamics at low physiological expression levels, limiting the number of photons that can be collected before the signal disappears ^1^.

Thus, the recent discovery of a new GFP variant from the cnidarian *Cytaeis uchidae* that compared with *Aequorea victoria* GFPs is several-fold brighter and more photostable could be a game changer for live microscopy approaches ^2^. However, this original StayGold is an obligate dimer limiting its usefulness for protein tagging. Motivated by this challenge, three different groups recently published very different versions of monomeric StayGold (Fig. 1a). Based on the crystal structure of the dimer, the original StayGold discoverers used targeted mutagenesis to isolate mStayGold, with nine amino acid substitutions mostly near the dimerization interface ^3^. Another group used random mutagenesis and screening to generate bright monomeric Staygold variants and isolated mBaoJin in which eight amino acid changes are more broadly distributed throughout the structure ^4^. Lastly, a third group identified a specific glutamate residue in the dimerization interface and generated a variant with a single E138D substitution to disrupt StayGold dimerization ^5^. This trifecta of monomeric StayGold variants was published within months of one another and raises the question which is the objectively brightest and most photostable variant to use for live microscopy protein tagging experiments as no independent side-by-side comparison of these variants has been made.

**Figure 1.**
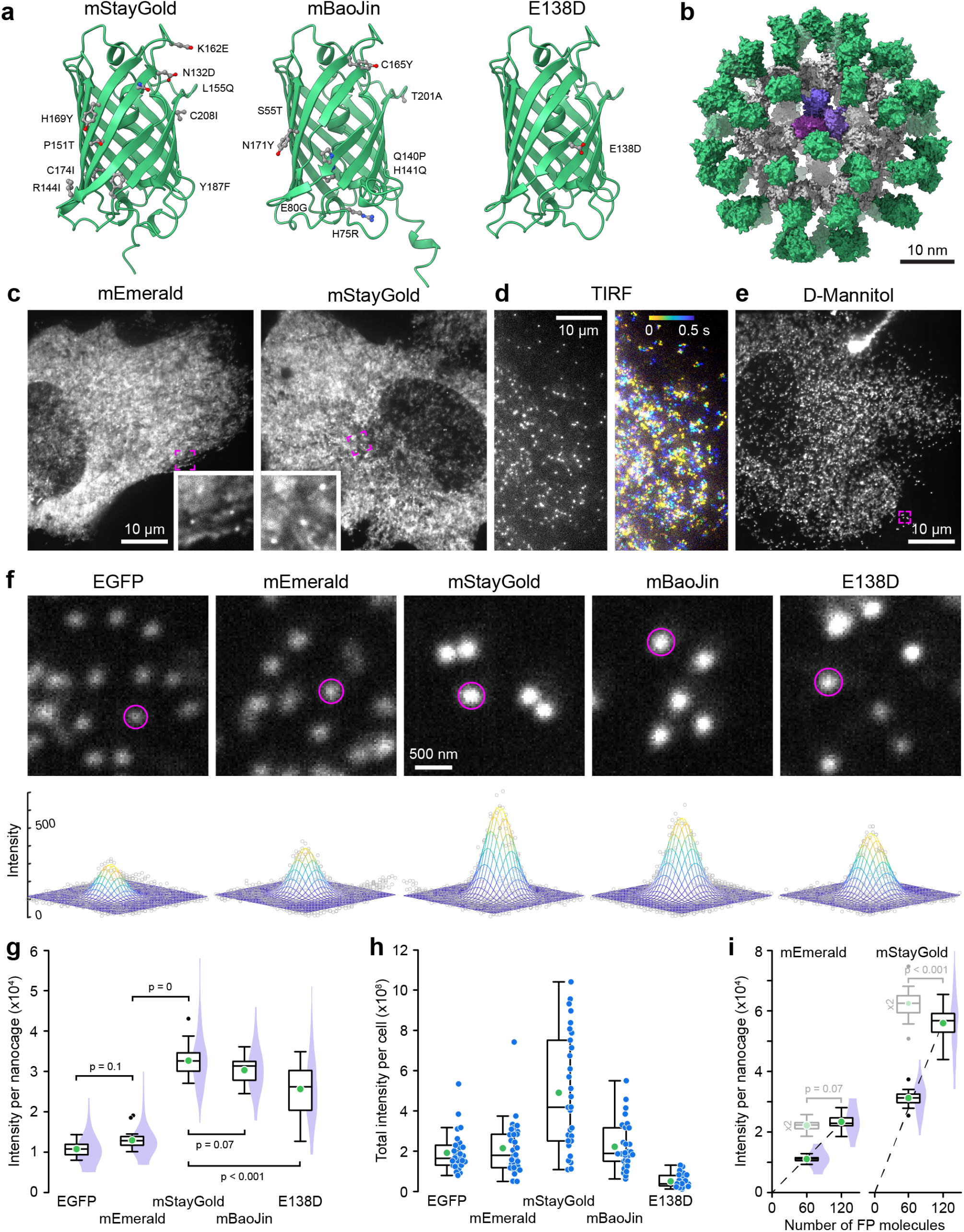
Comparison of FP-tagged I3-01 nanocage fluorescence intensity. (a) AlphaFold2 structures of the three monomeric mStayGold variants viewed from the dimerization interface. Amino acid substitutions compared to the original StayGold dimer are highlighted. (b) AlphaFold2 model of an mEmerald nanocage. The I3-01 peptides are grey with one trimer highlighted in shades of purple. (c) Spinning disk confocal images of mEmerald and mStayGold nanocage-expressing RPE cells at 500 ms exposure times. Insets show indicated regions at higher magnification highlighting immobile nanocages. (d) mStayGold nanocages in TIRF microscopy at 10 ms exposure times. The right panel shows a color-coded projection over 500 ms showing diffusion over this time period. (e) mStayGold nanocages by spinning disk confocal microscopy in a 400 mOsm D-Mannitol-treated RPE cell. The indicated region is shown at higher magnification in the middle panel below. (f) Comparable peripheral cell regions of D-Mannitol-treated RPE cells expressing the indicated FP nanocage fusions. Circles indicate areas within a two standard deviation radius as determined by 2D Gaussian fit. Bottom panels show the Gaussian fit for these example nanocage particles. (g) Comparison of the integrated brightness of nanocage particles tagged with the indicated FP. n = 30 cells each from 3 independent experiments. Statistical analysis by ANOVA with Tukey-Kramer HSD. (h) Total integrated intensity per cell of the same cell population as in (g). (i) Comparison of the integrated brightness of mEmerald and mStayGold nanocages containing either 60 or 120 FP molecules. The dashed lines are linear regressions through the origin. Box plots in light grey are the 60 FP molecule nanocage intensity data multiplied by two indicating the expected distribution if the 120 FP molecule nanocages were exactly twice as bright. Statistical comparison of these expected distributions to the 120 FP molecule nanocage intensities by Student’s t-test. n = 20 cells per condition. In (c), (e) and (f) all spinning disk confocal images were acquired with the same exposure settings and images are scaled to the same absolute intensities. In (g) - (i) green circles indicate the mean. Violin plots show the intensity distribution of all nanocage particles with 10 nanocages analyzed per cell.

To quantitatively compare monomeric StayGold variants to each other and to EGFP and mEmerald as reference standards that have been widely utilized in cell biology, we expressed FP-tagged I3-01 peptides in human retinal pigmental epithelial (RPE) cells. I3-01 is derived from trimeric aldolase ^6^, and sixty I3-01 subunits self-assemble into stable 26 nm diameter dodecahedron nanocages in which each of the vertices consists of one I3-01 trimer. We attached FPs to the I3-01 N-terminus because recent nanocage structures indicate that the N-terminus points toward the outside of the nanocage and is therefore less likely to interfere with nanocage assembly ^7^. Thus, each nanocage carries sixty FPs (Fig. 1b) allowing comparison of the absolute fluorescence intensity between individual nanocages with different FPs.

We imaged RPE cells expressing FP-tagged nanocages by spinning disk confocal microscopy because this modality is widely used for live microscopy. All spinning disk experiments used the same 500 ms exposure with a 488 nm irradiance of ∼15 W cm^-2^ to achieve high signal to noise. Although we were able to detect sub-resolution nanocage particles, most of the fluorescence signal was diffusely distributed throughout the cytoplasm (Fig. 1c). Since nanocages are expected to move freely within the cytoplasm, we hypothesized that this diffuse signal is caused by motion blur of nanocage particles due to the relatively long exposure time. Indeed, individual fast-moving nanocages were easily detected by TIRF microscopy at 50-times shorter exposure times (Fig. 1d). To slow intracellular diffusion, we therefore increased cytoplasm viscosity by treating cells with 400 mOsm D-Mannitol ^8^. Although this hypertonic treatment did not completely stop nanocage movement, it enabled observation of a sufficiently large number of nanocage particles at longer exposure times with very little background fluorescence (Fig. 1e, f). To compare the brightness of different FP-tagged nanocages, we fitted 2D Gaussian distributions to individual sub-resolution nanocage particles and integrated the fluorescence intensity inside a circle with a radius of two standard deviations. This direct comparison showed that all monomeric StayGold variant nanocages were significantly brighter than EGFP and mEmerald (Fig. 1g), which surprisingly were not significantly different from one another. mStayGold nanocages were approximately three times as bright as EGFP. mStayGold also appeared slightly brighter than mBaoJin, although this difference was not statistically significant. In contrast, E138D was significantly less bright. In addition, the fluorescence intensity distribution of the E138D nanocage particle population was highly variable indicating either incomplete nanocage assembly or aggregation (Fig. 1f). To compare expression levels in RPE cells, we measured the total brightness of transfected cells (Fig. 1h). Although a large variability is expected in transient transfections, on average mStayGold-expressing cells were more than 2.5-times brighter than EGFP-expressing cells indicating similar expression levels. In contrast, E138D-expressing cells were very dim possibly because this variant is not codon-optimized for mammalian expression. To test if high local concentrations affect fluorescence, we analyzed mEmerald and mStayGold nanocages with two FP tags in tandem. In contrast to mEmerald nanocages for which the fluorescence intensity increased linearly, nanocages with 120 mStayGold molecules were approximately 10% less bright than expected (Fig. 1i). This may indicate that mStayGold is more prone to fluorescence quenching at high local concentrations, which needs to be considered if nanocages are used as fluorescence standards in microscopy ^9,10^.

Photostability is the second critical performance parameter for live microscopy experiments. We therefore determined the photobleaching rate of these five FPs by measuring the decay in whole cell fluorescence under continuous 15 W cm^-2^ illumination (Fig. 2a-c). Determining the photobleaching rate constant by exponential fitting of each cell bleaching curve, we found that mStayGold was nearly three times as photostable (t_1/2_ = 60 +/-5.6 s; mean +/-standard deviation) as EGFP (t_1/2_ = 23 +/-1.2 s). To our surprise, however, mBaoJin photobleached rapidly with kinetics indistinguishable from EGFP (t_1/2_ = 20 +/-1.8 s).

**Figure 2.**
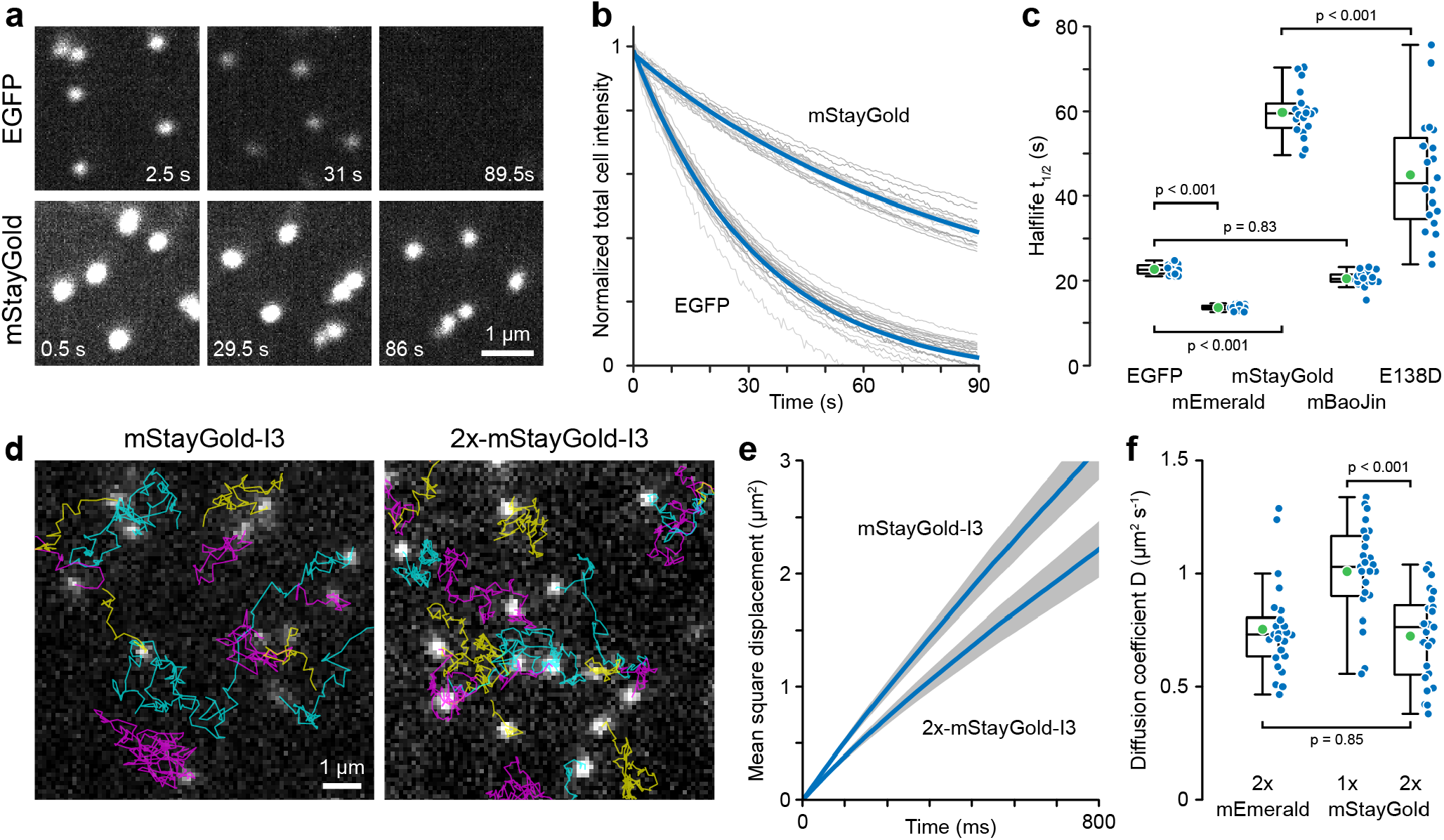
Comparison of FP-tagged I3-01 nanocage photostability and intracellular diffusion. (a) Representative EGFP and mStayGold nanocage photobleaching images. Images are scaled to the same absolute intensities. (b) Photobleaching as a function of light exposure. Grey curves are measurements from individual cells normalized to the fluorescence intensity at the beginning of the time series. Blue curves are single exponential fits. (c) Comparison of photobleaching half-life. n = 20 cells per condition. Statistical analysis by ANOVA with Tukey-Kramer HSD. (d) Representative TIRF microscopy images from 85 frames per second time series to measure nanocage diffusion. Particle tracks are overlaid in color. (e) Average mean square displacement curves from 25 cells per condition representing a total of 16504 mStayGold and 20692 2x-mStayGold nanocage particle tracks. Grey areas indicate 95% confidence intervals. (f) Comparison of the diffusion coefficients of the indicated nanocage constructs. n = 25 cells per condition. Statistical analysis by ANOVA with Tukey-Kramer HSD.

Although all FP-tagged nanocages appeared as sub-diffraction objects with a full width at half maximum (FWHM) near the resolution limit of our spinning disk microscope system (mEmerald: 202 +/-20 nm; mStayGold: 208 +/-20 nm), to ascertain that the increased brightness of mStayGold nanocages was not due to aggregation, we compared intracellular diffusion of mEmerald and mStayGold-tagged nanocages. Nanocage movement was tracked in image sequences acquired by TIRF microscopy at 85 frames per second (Fig. 2d) and apparent diffusion coefficients determined by mean square displacement over time assuming Brownian motion (Fig. 2e). Nanocages with one mStayGold molecule per I3-01 peptide had an apparent diffusion coefficient of D = 1.01 +/-0.21 µm^2^ s^-1^ while the diffusion coefficient of 2x-mStayGold nanocages was reduced to 0.72 +/-0.20 µm^2^ s^-1^. At low Reynolds number, the diffusion coefficient is approximately inversely linearly related to particle size and this ∼25% decrease in diffusion coefficient is consistent with a ∼25% increase in effective nanocage size due to addition of another layer of mStayGold molecules to the nanocage exterior (Fig. 1b). Importantly, the diffusion coefficient of 2x-mEmerald nanocages (D = 0.76 +/-0.20 µm^2^) was indistinguishable from 2x-mStayGold nanocages indicating that these particles have the same size and therefore mStayGold nanocages do not aggregate (Fig. 2f). These diffusion coefficients are also in a similar range as cytoplasm diffusion coefficients measured for similar sized nanoparticles ^11,12^.

In conclusion, we present a new method to compare fluorescent proteins on a molecule-by-molecule basis in physiological conditions independent of relative expression levels or ratio measurements to other fluorescent proteins. Our method is simple and can be easily adopted to other imaging modalities or cell types to determine which FP is optimal for a specific condition. In mammalian cells, our analysis reveals a clear favorite. Although mStayGold and mBaoJin are both much brighter than EGFP, in practical terms the exceptional mStayGold photostability combined with the increased brightness allows nearly ten times as many exposures as EGFP with a similar fluorescence signal at lower illumination power. Our diffusion measurements further show that mStayGold nanocages can serve as improved monodisperse photostable nanoparticles for the biophysical characterization of intracellular environments.

## Online Methods

### Molecular Cloning

I3-01 nanocage constructs tagged either with a single or a tandem repeat of the indicated fluorescent proteins at the N-terminus were constructed by Gibson assembly with 4-7 GGS spacers between the FP and the I3-01 nanocage peptide. The mStayGold coding sequence was PCR amplified from pCMV3-3xNLS-mStayGold (a gift from Cuyler Luck). The mBaoJin coding sequence was synthesized as a gene fragment (TWIST Biosciences) based on the sequence reported in ^4^. The E138D coding sequence was PCR amplified from pcDNA3-10His-mStayGold(E138D) which was a gift from Mohan Balasubramanian (Addgene plasmid #211363; RRID:Addgene_211363). All constructs were verified by whole plasmid sequencing (Primordium Labs). Protein structural models were computed using the AlphaFold2 Google Colab ^13^.

### Cell culture and live microscopy

RPE cells were cultured in DMEM/F12 medium (Invitrogen) supplemented with 10% fetal bovine serum (Atlanta Biosciences) and 2 mM glutamine (Invitrogen) at 37°C, 5% CO^2^. For transient nanocage expression and microscopy, cells were plated in 35-mm #1.5 glass-bottom dishes (MatTek), transfected after 24 h using jetOPTIMUS (Sartorius), and used for experiments 24 h later. To increase extracellular osmolarity, the tissue culture medium was exchanged with medium supplemented with 0.28 mM (i.e. 400 mOsm) D-Mannitol 10-15 min before imaging.

High-resolution spinning disk image sequences of nanocage-expressing RPE cells were acquired with a CFI Apochromat TIRF 60x NA 1.49 CFI Apochromat objective (Nikon) on a Yokogawa CSU-W1/SoRa spinning disk confocal system and an ORCA Fusion BT sCMOS camera (Hamamatsu) controlled through NIS Elements v5.3 software (Nikon) yielding an effective pixel size of 27 nm. TIRF image sequences were acquired on a TI inverted microscope (Nikon) equipped with a motorized TIRF illuminator (Nikon) with a 100x 1.49 NA CFI Apochromat TIRF objective (Nikon) using 1.5x intermediate magnification and an iXon EMCCD camera (Andor) yielding an effective pixel size of 110 nm. The camera ROI was set to 150 × 512 pixels to achieve 85 frames per second acquisition rates.

### Image analysis

The integrated fluorescence intensity of individual nanocages was quantified with custom-written MATLAB code to fit 2D Gaussian functions to user-selected nanocage particles. Fluorescence intensity from the original image data inside a circle with a radius of two standard deviations, which accounts for approximately 86% of the total volume of a 2D Gaussian distribution ^14^, around the maximum of the Gaussian fit minus the offset of the fit representing the local background was integrated to calculate the nanocage fluorescence signal. To analyze photobleaching, the average intensity in an ROI encompassing the entire cell was measured over time and fitted with an exponential decay function to determine photobleaching rate constants.

Nanocage particles in TIRF image sequences were tracked in MATLAB using the u-track package (https://github.com/DanuserLab/u-track)^15^ with Gaussian mixture model detection and default settings. Individual nanocage particle tracks longer than 20 frames and shorter than 250 frames (i.e. <3 s) were extracted from the ‘tracksFinal’ structure, and diffusion coefficients determined by linear fitting of the initial 25% of the average mean square displacement curve per cell using the @msdanalyzer package (https://tinevez.github.io/msdanalyzer/)^16^ assuming Brownian motion at short time scales.

### Statistics

Details of statistical analysis including the type of test, p-values and numbers of biological replicates are provided within the relevant figures and figure legends. All statistical analysis was done in MATLAB (Mathworks, Inc.)., and graphs were produced in MATLAB and in Excel (Microsoft), and figures assembled with Adobe Illustrator. In all figures, box plots show median, first and third quartile, with whiskers extending to observations within 1.5 times the interquartile range.

## Acknowledgements

We thank Cuyler Luck and Mohan Balasubramanian for StayGold plasmids. This work was supported by National Institutes of Health grants R61 CA278518, S10 RR026758 and S10 OD028611 to T.W. K.A.J. is supported by Tobacco-Related Disease Research Program Predoctoral Fellowship T33DT6442. This material is based upon work supported by the National Science Foundation Graduate Research Fellowship Program under Grant No. 2038436 (to K.A.J.). Any opinions, findings, and conclusions or recommendations expressed in this material are those of the author(s) and do not necessarily reflect the views of the National Science Foundation.

## Supplementary Videos

**Videos 1.** mStayGold I3-01 nanocage diffusion by TIRF microscopy in transfected RPE cells. Acquisition at 22 frames s^-1^.

**Videos 1.** Photobleaching comparison of EGFP and mStayGold I3-01 nanocages in 400 mOsm D-mannitol-treated RPE cells by spinning disk confocal microscopy under continuous 15 W cm^-2^ illumination. Images are scaled to the same absolute intensities and shown in pseudocolor. Note that the mStayGold nanocages at the end of the video are still brighter than the EGFP nanocages in the beginning. Acquisition at 2 frames s^-1^.

